# Mixology: a tool for calculating required masses and volumes for laboratory solutions

**DOI:** 10.1101/2021.01.30.428948

**Authors:** Theo Sanderson

## Abstract

We have created a tool to calculate the amount of each ingredient required to make up a custom solution with defined concentrations. It can convert between many kinds of volumetric, mass and concentration units. This includes the ability to convert between molarities and mass-based concentrations, using molecular masses retrieved from the ChEBI database. Mixology can be accessed at http://mixology.science.

## Introduction

Performing calculations to make up solutions is one of the first skills that a laboratory researcher learns. Despite efforts to facilitate sharing of laboratory protocols (1), solutions are often described in publications only in terms of the molarities of their constituent parts, meaning that calculations are needed to determine the absolute amounts needed to make up some defined volume. These calculations can be surprisingly lengthy.

Suppose a researcher is implementing a new protocol, and needs to make a buffer containing, amongst other ingredients, “500 μg/ml sodium chloride”. She wants to make up 2 liters of the new buffer. On her bench she happens to already have a stock solution labelled “1.5 M NaCl”. What volume of stock solution does she need to add for the new buffer? One approach that she might follow to calculate this would be to:

1. Look up the molar mass of sodium chloride:

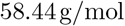
2. Convert the desired concentration, 500 μg/ml, to μg^/L:^

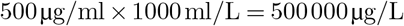
3. Convert the desired concentration to g/L:

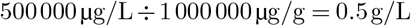
4. Convert the desired concentration to mol/L (molarity):

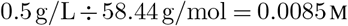
5. Calculate the ratio of the two concentrations:

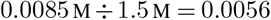
6. Calculate the volume to add:

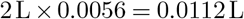
7. Convert this volume to ml for pipetting:

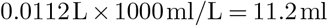

If her buffer contains five different such ingredients, she might need to perform a similar set of steps five times, requiring perhaps 35 individual steps. If an error occurs in just one of these, the final buffer will not be to the expected specification.

We decided to build a simple tool to perform calculations such as these (Fig. 1). Mixology automates the retrieval of molecular masses, and the mathematical operations needed to calculate the required masses of reagent, or volumes of stock solutions, needed for desired final concentrations. It is available at http://mixology.science.

**Fig. 1.**
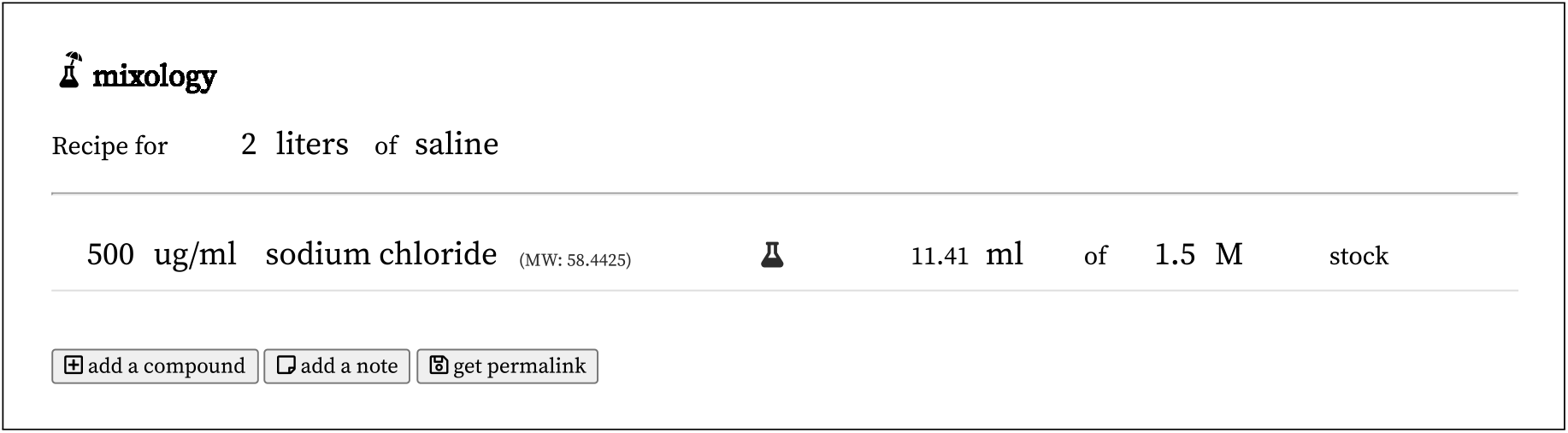
A screenshot of the Mixology interface, featuring the single compound solution described in the Introduction. The permalink for this recipe is https://mixology.science/?recipe=ngEh5UbDZX.

**Fig. 2.**
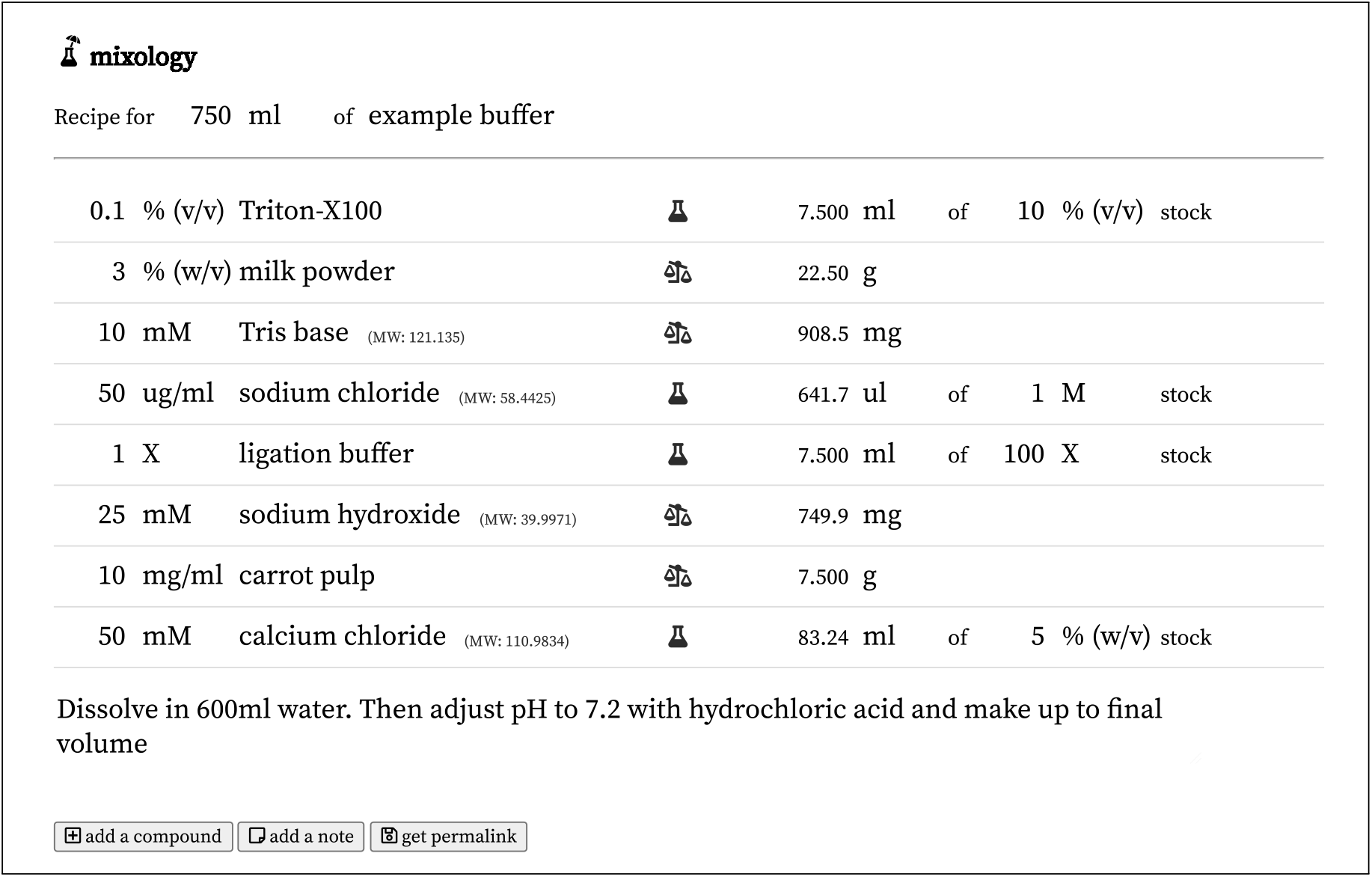
A screenshot of the Mixology interface, with an example buffer demonstrating many of the different classes of interconversion of which Mixology is capable. The permalink for this recipe is https://mixology.science/?recipe=mN1j1sNSD5.

### Usage

Mixology provides a simple interface to record the specifications of a solution and calculate the required amount of each ingredient (Fig. 1). A user first enters the volume of the solution they want to make, and a descriptive name. Then they add any number of components to the solution. For each component, they enter a desired final concentration. Concentrations can be entered in any of several unit forms:

- mass-based: g/ml, μg/ml, mg/l, % (w/v), etc.
- molarity-based: mol/l, nM, μM, mM, M, etc.
- volume-based: % (v/v), X
- other: units/ml, etc.

The user also enters the name of the chemical. Chemical names are autocompleted with known entities from ChEBI (2). Where a known chemical is entered, its molecular mass is inferred from the ChEBI database. Alternatively a custom compound name can be entered, and a molecular mass (if needed) entered manually.

If the desired concentration is mass- or molarity-based, the user can select whether to weigh out the compound (mass units) or to measure out a volume of stock solution (with a mass- or molarity-based stock concentration). If the desired concentration is volumetric, a volume of stock solution (which can be 100% (v/v)) is required.

In either case, the user enters the units of mass or volume to weigh out, and Mixology automatically calculates the quantity of this unit required to make up the final volume at the desired concentration.

The user can add as many components as required, and also add notes, which might contain instructions such as “adjust pH to 7.2”. Finally they can create a ‘permalink’ pointing to their solution, creating a unique URL for the solution that can be saved and shared.

### Implementation

The code behind Mixology is available in our GitHub repository (https://github.com/theosanderson/mixology).

Mixology is developed with the Vue.js framework. This allows reactive behaviour in which all fields are updated in real time as others are changed. We use the vue-simple-suggest component (3) to list available units and chemical entities from ChEBI.

All concentration terms are converted into a standard format consisting of some quantity (moles, grams, liters, activity units) per liter, facilitating downstream steps. Mol/l and g/l can be interconverted using the molecular mass, but these cannot be interconverted to the other unit types (volumetric and activity units).

Molecular masses, and compound names for autocompletion, are taken from ChEBI(2). ChEBI entities are processed into JSON with a custom R notebook, available in our GitHub repository, using R and the tidyverse (4, 5). We take as input ChEBI’s three-star chemicals, which are those that have been curated. We exclude any name that is mapped to multiple distinct chemicals to avoid potential confusion. Sometimes multiple mass measurements are available for a chemical in ChEBI. Normally these are extremely similar, so we exclude only chemicals for which the standard deviation of masses is greater than 0.001.

Recipes can be saved using a “permalink” feature. This deposits a copy of the recipe in JSON format into an external Firebase database, with a unique identifier which can be used to retrieve it at the correct URL.

## Discussion

We have created a simple tool which we hope will be useful to laboratory scientists. In the day following Mixology’s release on Twitter, a user testing it identified an inconsistency with the calculations in a published protocol (6). This turned out not to be a problem with the tool, but the use of an incorrect molar mass (hydrated vs. anhydrate) for calculations in the official protocol. This provides one example of how easily small mistakes can occur performing these operations manually.

Mixology has some limitations. It cannot currently perform any calculations which would require knowing the density of a substance (interconversion between mass and purely volumetric units). Additionally, it does not provide the facility to calculate quantities required to obtain a particular pH or osmolarity. We welcome feature suggestions and pull requests in our GitHub repository.

We do not argue that the availability of a tool such as this replaces the need for scientists to know how to perform these types of calculations. However, we hope that by providing a streamlined tool that eases this process, Mixology can make one aspect of routine laboratory life simpler, and reduce errors.

## Acknowledgements

This manuscript uses a LaTeX template developed by Ricardo Henriques.

## Notes

http://mixology.science

https://github.com/theosanderson/mixology

### Competing Interest Statement

The authors have declared no competing interest.

